# Evaluating the granularity and statistical structure of lesions and behaviour in post-stroke aphasia

**DOI:** 10.1101/802595

**Authors:** Ying Zhao, Ajay D. Halai, Matthew A. Lambon Ralph

## Abstract

The pursuit of relating the location of neural damage to the pattern of acquired language and general cognitive deficits post-stroke stems back to 19^th^ century behavioural neurology. Whilst spatial specificity has improved dramatically over time, from the large areas of damage specified by post-mortem investigation to the millimetre precision of modern MRI, there is an underlying issue that is rarely addressed, which relates to the fact that damage to a given area of the brain is not random but constrained by the brain’s vasculature. Accordingly, the aim of this study was to uncover the statistical structure underlying the lesion profile in chronic aphasia post-stroke. By applying varimax-rotated principal component analysis to the lesions of 70 patients with chronic post-stroke aphasia, we identified 17 interpretable clusters, largely reflecting the vascular supply of middle cerebral artery sub-branches and other sources of individual variation in vascular supply as shown in classical angiography studies. This vascular parcellation produced smaller displacement error in simulated lesion-symptom analysis compared with individual voxels and Brodmann regions. A second principal component analysis of the patients’ detailed neuropsychological data revealed a four factor solution reflecting phonological, semantic, executive-demand and speech fluency abilities. As a preliminary exploration, stepwise regression was used to relate behavioural factor scores to the lesion principal components. Phonological ability was related to two components, which covered the posterior temporal region including the posterior segment of the arcuate fasciculus, and the inferior frontal gyrus. Three components were linked to semantic ability and were located in the white matter underlying the anterior temporal lobe, the supramarginal gyrus and angular gyrus. Executive-demand related to two components covering the dorsal edge of the middle cerebral artery territory, while speech fluency was linked to two components that were located in the middle frontal gyrus, precentral gyrus, and subcortical regions (putamen and thalamus). Future studies can explore in formal terms the utility of these PCA-derived lesion components for relating post-stroke lesions and symptoms.

## Introduction

Classically, investigators used post mortem dissections to provide insight into which brain areas were related to different behaviours/functions. These included the seminal studies of Broca and Wernicke who identified areas of the brain related to speech production and comprehension, respectively (Broca, 1861; Wernicke, 1874). A subsequent development allowed the general area of damage to be explored, *in vivo*: one consequence of the tragic World Wars was that, for soldiers who survived missile head injuries, the trajectory of the missile could be determined (by entry/exit points). This allowed researchers to infer the location of damage and determine the effects on behaviour in greater detail (e.g. Head, 1926; Schiller, 1947; Tallal & Newcombe, 1978). Contemporary *in vivo* lesion mapping became possible with the invention of CAT (Ambrose & Hounsfield, 1973) and MRI (Mansfield & Maudsley, 1977). MRI scanners can now image tissue with great precision (< 1mm^3^) and novel acquisition protocols allow imaging of different tissue properties (in structural and functional modalities). This engineering technology has been combined with advances in analytical techniques (Bates et al., 2003; Tyler, Marslen-Wilson, & Stamatakis, 2005) resulting in increasingly sophisticated, detailed lesion-symptom mapping (though subject to significant challenges: cf. Mah et al., 2014) and the evaluation of lesion-based prediction models.

Lesion mapping methodologies have been used widely in the stroke aphasia literature (e.g. Bates et al., 2003; Geva et al., 2011; Magnusdottir et al., 2013; Mirman, Chen, et al., 2015b; Schwartz et al., 2009; Yourganov, Fridriksson, Rorden, Gleichgerrcht, & Bonilha, 2016). While the precision of the structural imaging has been improved dramatically, there is an underlying issue that is rarely addressed: damage to a given area of the brain after stroke is not random but constrained by the neurovasculature. This means that there is neither full nor random sampling of the brain, which would be the ideal situation for lesion-symptom mapping. In addition, the non-random nature of stroke means that there is some degree of co-linearity between neighbouring voxels, which can bias or mislocalise lesion-symptom relationships (Inoue, Madhyastha, Rudrauf, Mehta, & Grabowski, 2014; Mah, Husain, Rees, & Nachev, 2014). Classical angiography studies identified the regions supplied by the arterial branches as well as their variations across individuals (Michotey, Moscow, & Salamon, 1974; Newton & Potts, 1974). For the cognitive and language deficits associated with MCA stroke, the most pertinent branches are probably those related to the subcortical regions (arising from M1), the insular (arising at M2) and the eleven cortical branches (arising from M1 and M4), including: 1) orbitofrontal, 2) prefrontal, 3) precentral, 4) central, 5) anterior parietal, 6) posterior parietal, 7) angular, 8) temporo-occipital, 9) posterior temporal, 10) middle, temporal, 11) anterior and polar temporal arteries. Accordingly, voxels that fall into each of these specific regions are likely to be damaged together and, in turn, there will be correlations between certain cortical regions due to the bifurcation/trifurcation of the MCA (e.g., occlusion of the superior branch will affect prefrontal and motor regions but not those of the temporal lobe).

The key aim of this study, therefore, was to use a data-driven approach to uncover the underlying patterns in the lesions of seventy chronic aphasic post-stroke cases. We achieved this aim by using principal component analysis (PCA) with varimax rotation to investigate the patients’ lesion structure. One previous study on coma used PCA on brain images to compare with a probabilistic frequency map (Singhal et al., 2012). They identified six components in the patients’ data and concluded that the PCA methodology was better equipped to depict patterns of co-varying damage than probabilistic frequency maps. In addition, we tested whether the new parcellation could reduce the displacement error observed in single voxel lesion-symptom mapping by replicating a pipeline developed in a recent study (Mah et al., 2014). We compared the lesion-based parcellations and voxel-based results to another widely-used anatomical parcellation (Brodmann areas) as a control.

Finally, we also conducted an initial exploratory analysis to explore how the data-driven lesion-based parcellations relate to the patients’ behavioural variations. In previous work, multiple research groups have used PCA with varimax rotation to unpack complex behavioural variations (Butler, Lambon Ralph, & Woollams, 2014; Corbetta et al., 2015; Fucetola, Connor, Strube, & Corbetta, 2009; Halai, Woollams, & Lambon Ralph, 2017; Kümmerer et al., 2013; Lacey, Skipper-Kallal, Xing, Fama, & Turkeltaub, 2017; Mirman, Chen, et al., 2015b; Nair et al., 2015; Siegel et al., 2016; Vandenbulcke, Peeters, Van Hecke, & Vandenberghe, 2005; Wilson et al., 2010) The main advantage of applying varimax rotation is to improve interpretability by rotating the principal components such that assessments load highly on one component and minimally on others (Jolliffe, 2002). When applied to data collected from a large group of chronic, post-stroke aphasics across a set of 21 detailed language and cognitive assessments, we previously identified four factors that included phonological ability, semantic ability, executive-demand and speech fluency (Butler et al., 2014; Halai et al., 2017). Similar data structures have been obtained in independent patient samples (Alyahya, Halai, Conroy, & Lambon Ralph, 2018; Halai, Woollams, & Lambon Ralph, 2018; Kümmerer et al., 2013; Mirman, Chen, et al., 2015a; Mirman, Zhang, Wang, Coslett, & Schwartz, 2015). This study utilised the same deconstruction of the patients’ behavioural data and compared two different forms of exploratory lesion-symptom analyses: (1) the more traditional approach based on independent voxel analyses; and (2) an investigation of the potential relationship between the behavioural and lesion PCA structures.

## Materials and methods

### Participants

This study included 70 post-stroke patients with chronic aphasia (either ischaemic or haemorrhagic) with damage restricted to the left hemisphere (see Supplementary Table 1 for patients’ background information). Inclusion criterion for recruiting patients were: (1) monolingual native English speakers, (2) normal or corrected-to-normal hearing and vision, (3) right-handed, (4) one stroke, (5) at least 12 months’ post-stroke, (6) no other known neurological conditions, (7) no contradistinctions for MRI scanning and (8) chronic aphasia of any type or severity. Informed consent was obtained from all participants under approval from the local ethics committee. Structural imaging data from a healthy age and education matched control group (8 female, 11 male) was used to determine the lesion outline in the patients using an automated lesion identification procedure (Seghier, Ramlackhansingh, Crinion, Leff, & Price, 2008). We note that a subset of patients (N = 31) was used in previous studies (Butler et al., 2014; Halai et al., 2017).

**Table 1.** Loadings of behavioural assessments on factors extracted from the rotated PCA.

| Tasks | Component |  |  |  |
| --- | --- | --- | --- | --- |
|  | Phonology | Semantics | Executive | Fluency |
| Delayed Repetition - Words | **0.888** | 0.221 | 0.183 | 0.193 |
| Delayed Repetition - Non-words | **0.883** | 0.027 | 0.237 | 0.148 |
| Immediate Repetition - Non-words | **0.881** | 0.061 | 0.231 | 0.143 |
| Immediate Repetition - Words | **0.858** | 0.211 | 0.125 | 0.170 |
| Boston Naming Test | **0.823** | 0.381 | 0.077 | 0.121 |
| 64-Item Naming | **0.813** | 0.431 | 0.154 | 0.117 |
| Forward Digit Span | **0.746** | 0.233 | 0.188 | 0.073 |
| Backward Digit Span | **0.595** | 0.207 | 0.234 | 0.359 |
| CAT Spoken Sentence Comprehension | **0.521** | 0.455 | 0.441 | 0.163 |
| Spoken Word to Picture Matching | 0.236 | **0.801** | 0.267 | 0.145 |
| Type/Token ratio | 0.362 | **0.718** | -0.075 | -0.092 |
| Written Word to Picture Matching | 0.182 | **0.713** | **0.504** | 0.155 |
| Camel and Cactus Test: Pictures | 0.092 | **0.688** | 0.484 | 0.288 |
| 96 Synonym Judgement | 0.381 | **0.658** | 0.315 | 0.359 |
| Minimal Pairs - Non-words | 0.353 | 0.058 | **0.814** | -0.014 |
| Raven's Coloured Progressive Matrices | 0.048 | 0.274 | **0.735** | 0.156 |
| Minimal Pairs - Words | 0.419 | 0.168 | **0.705** | 0.132 |
| Brixton Spatial Anticipation Test | 0.132 | 0.178 | **0.698** | 0.231 |
| Token | 0.010 | 0.034 | 0.207 | **0.885** |
| Mean Length of Utterance in Morphemes | 0.314 | 0.252 | 0.137 | **0.831** |
| Words-Per-Minute | 0.314 | 0.096 | 0.080 | **0.768** |

Factor loadings > 0.5 are given in bold; CAT = Comprehensive Aphasia Test.

### Neuropsychological assessments

To test the participants’ speech, language and cognitive abilities, we utilised a detailed neuropsychological test battery, designed to assess input/output phonological processing, semantic processing and sentence comprehension, as well as general executive-cognitive function. The battery included a subset of tasks from the Psycholinguistic Assessments of Language Processing in Aphasia (PALPA) battery (Kay, Lesser, & Coltheart, 1992): (1) auditory discrimination using non-word minimal pairs, (2) auditory discrimination using word minimal pairs, (3) immediate repetition of non-words, (4) immediate repetition of words, (5) delayed repetition of non-words, and (6) delayed repetition of words. Tasks from the 64-item Cambridge Semantic Battery (Bozeat, Lambon Ralph, Patterson, Garrard, & Hodges, 2000) were included: (7) word-to-picture matching task (spoken version), (8) word-to-picture matching task (written version), (9) Camel and Cactus Test (picture), and (10) picture naming test. Other language tasks included (11) the Boston Naming Test (BNT) (Kaplan, Goodglass, & Weintraub, 1983), (12) written 96-trial synonym judgement test (Jefferies, Patterson, Jones, & Lambon Ralph, 2009), (13) the spoken sentence comprehension task from the Comprehensive Aphasia Test (CAT) (Swinburn, Baker, & Howard, 2005), and the ‘Cookie theft’ picture description task from the Boston Diagnostic Aphasia Examination (Goodglass & Kaplan, 1983). Specifically, patients’ responses in the ‘Cookie theft’ picture description task were recorded and transcribed. The (14) number of word tokens (T), (15) type/token ratio (TTR), (16) mean length of utterance in morphemes (MLU), and (17) words-per-minute (WPM) were computed. Cognitively related tasks included (18) forward and (19) backward digit span (Wechsler, 1987), (20) the Brixton Spatial Rule Anticipation Task (Burgess & Shallice, 1997), and (21) Raven’s Coloured Progressive Matrices (Raven, 1962). All scores were converted into percentage based on the maximum score available; where no maximum was available, we used the max score in the group. The testing took place over a number of testing sessions, where the pace was determined by the participant and each session lasted approximately 1 hour and 30 minutes. The testing pipeline was as follows: 1) initial consent and start testing battery, 2) MRI scan and 3) continue behavioural testing to completion, where all testing was completed within two months of starting the testing.

### Acquisition of Neuroimaging Data

High resolution structural T1-weighted MRI scans were acquired on a 3.0 Tesla Philips Achieva scanner (Philips Healthcare, Best, The Netherlands) using an 8-element SENSE head coil. A T1-weighted inversion recovery sequence with 3D acquisition was employed, with the following parameters: TR (repetition time) = 9.0 ms, TE (echo time) = 3.93 ms, flip angle = 8°, 150 contiguous slices, slice thickness = 1 mm, acquired voxel size 1.0 × 1.0 × 1.0 mm^3^, matrix size 256 × 256, FOV = 256 mm, TI (inversion time) = 1150 ms, SENSE acceleration factor 2.5, total scan acquisition time = 575 s.

### Preprocessing Neuroimaging Data

High-resolution structural scans were pre-processed with the same procedure as our previous studies (Butler et al., 2014; Halai et al., 2017), with one additional first step. We extracted the brain space for all T1 images using a brain extraction tool optimised for patient brains (Lutkenhoff et al., 2014). This step was performed in order to avoid edge artefacts during smoothing where non-brain tissue with high intensity would be smoothed into brain tissue. We obtained a mask in native space and used the following procedure to normalise the whole T1 image (not brain extracted) and use the transformation matrix to warp the mask into MNI space. The analysis used Statistical Parametric Mapping software (SPM8: Wellcome Trust Centre for Neuroimaging, http://www.fil.ion.ucl.ac.uk/spm/) and a modified unified segmentation-normalisation procedure (Seghier et al., 2008). After normalising individual lesioned brain images into standard Montreal Neurological Institute (MNI) space, images were smoothed with an 8mm full-width-half-maximum (FWHM) Gaussian kernel. Lesions were automatically identified for each patient by comparing the structural image with the age and education matched control group, using an outlier detection algorithm to identify ‘abnormal’ voxels (Seghier et al., 2008). All parameters were kept at default except the lesion definition ‘U-threshold’, which was set to 0.5 after comparing the results obtained from a sample of patients to what would be nominated as lesioned tissue by an expert neurologist. The method produces two types of outputs: an abnormality likelihood map and a binary lesion map. In the abnormality likelihood map, the signal intensity for any given voxel is compared to the distribution of signal in the same voxel from a control group where a higher value indicates that the voxel is more likely to be abnormal (scaled 0-1). The binary lesion maps where defined using the ‘U-threshold’ and a cluster extent of 100 voxels (i.e., in our case the voxels had to have at least 50% likelihood of being abnormal and form clusters larger than 100 voxels). The abnormality likelihood maps were used in the PCA and the binary lesion maps were only used to show the distribution of lesions within our dataset (Figure 1).

**Figure 1.**
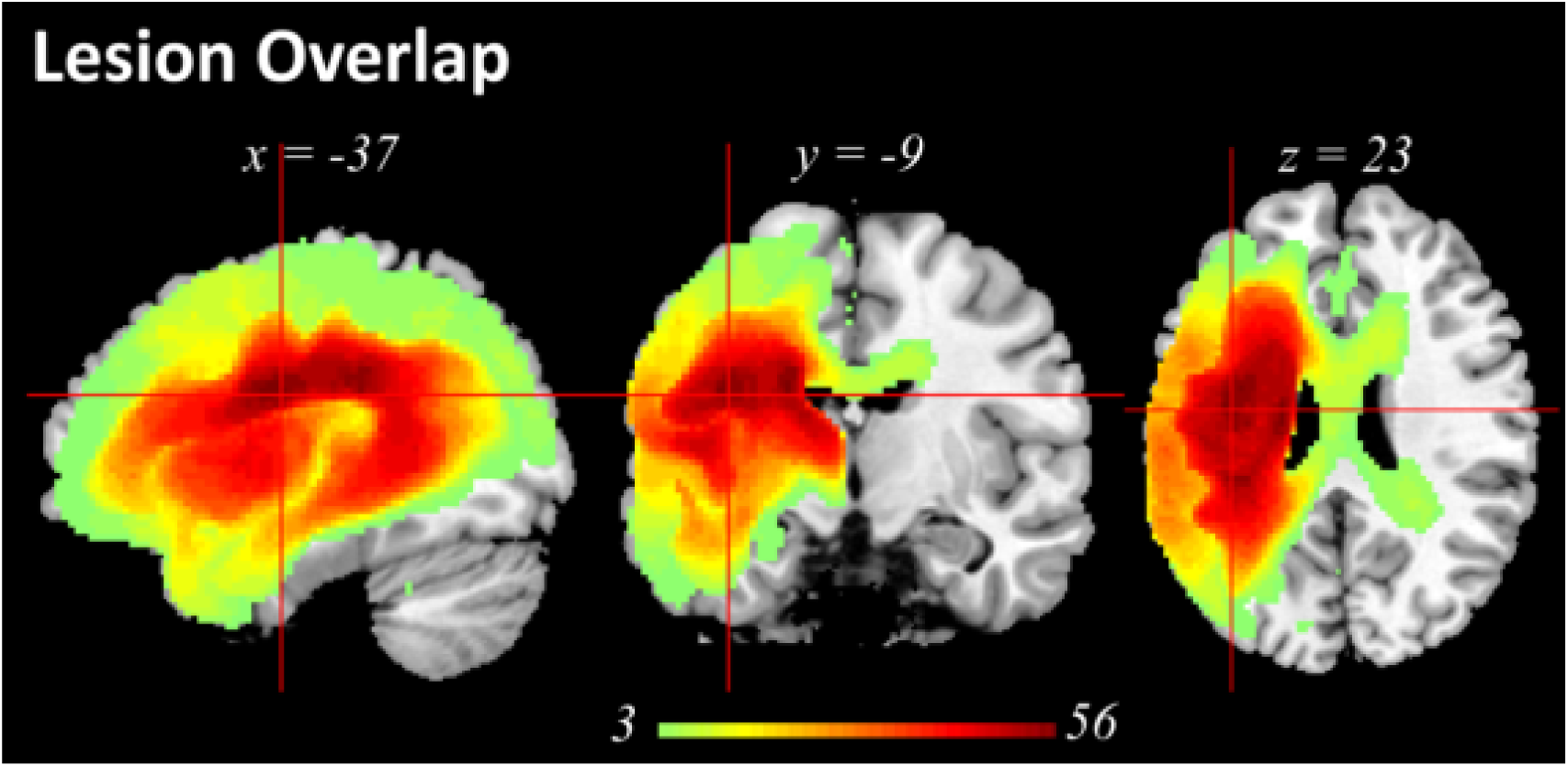
lesion profile of the patient group. Subcortical regions and white matter, e.g., putamen, corpus callosum, thalamus, inferior occipito-frontal fasciculus, caudate, internal capsule, insular, etc., had the largest probability of being damaged.

### Preliminary analyses of voxel-wise lesion to behavioural variations

We performed a principal component analysis using SPSS 22. The input data consisted of twenty-one behavioural scores for seventy patients. Factors were extracted based on the correlation matrix and components with eigenvalues greater than 1.0 were varimax rotated. Individual patient scores for each factor were obtained using the regression method. These factor scores were then entered into the same model (as they are orthogonal) and correlated to the lesion profile based on T1-weighted images at the voxel wise level (as done in Butler et al., 2014; Halai et al., 2017; Lacey et al., 2017; Mirman, Zhang, et al., 2015). Clusters were significant at AlphaSim corrected *p* < 0.01 with a voxel threshold of *p* < 0.001.

### Preliminary analyses of the relationships between PCA-based lesion components and behavioural variations

We conducted the PCA of the lesion data in MATLAB (R2012a) and used the minimum description length criteria and Kolmogorov information criterion (Akaike, 1974; Rissanen, 1983) (codes from GIFT Matlab software, http://icatb.sourceforge.net/) to estimate the number of components within the data that explained the most variance with a penalty for increasing the number of components. We used the “princomp” function to extract the components and then applied a varimax rotation to help with the interpretation of the resulting components (using the “rotatefactors” function). We projected the PCA factor scores back into brain space and converted the data into z-scores and used |*z*/ > 3.29 (*p* < 0.001 two-tail) as the cut-off value. Clusters larger than 500 voxels (volume: 4 cm^3^) were reported using the Harvard-Oxford (cortical and subcortical) and natbrainlab atlases (white matter). Having obtained the structure of the underlying lesions, we explored which lesion component(s) were related to the behavioural variations. First, we conducted Pearson correlation analysis between patients’ twenty lesion component scores and four behaviour component scores (Bonferroni corrected at *p* < 0.05). To test which of the lesion components were the most important predictors, we built a stepwise regression model where the lesion component scores were used to predict the behavioural component scores. The lesion component(s) that significantly predicted behavioural factors were then mapped onto the brain and converted into z-scores [|*z*/ > 3.29 (*p* < 0.001 two-tail) as the cut-off value]. Clusters larger than 200 voxels (volume: 1.6 cm^3^) were reported using the Harvard-Oxford (cortical and subcortical regions) and natbrainlab atlases (white matter).

### Assessing displacement error using different lesion models

We determined the displacement error induced by different lesion models (single voxel, lesion territory parcellation and Brodmann parcellation) in lesion-symptom mapping replicating the pipeline proposed by Mah and colleagues (2014). In the first instance, we simulated the ground truth at the voxel level, meaning that one single voxel is critical to a certain behaviour. For this voxel, we split our sample into two groups based whether or not the voxel was damaged and performed mass-univariate analyses to search for other voxels whose damage was associated to the ground truth. A Fisher’s exact test was used to determine statistical inference, where the resultant map was thresholded at *p* < 0.01, FDR corrected for multiple comparisons. We calculated the centre of mass of the significant cluster and determined the Euclidean distance between this point and ground truth voxel. This process was repeated for 57632 voxels, for which more than three patients had damage. Next, we simulated the ground truth using Brodmann areas, where for each area the sample was split into intact/damaged based on at least 20% of the area being damaged and a case with damage had a 90% chance of exhibiting a behavioural deficit (Mah et al., 2014). The same lesion mapping method was used as in the single voxel pipeline (Fisher’s exact test, p < 0.01, FDR corrected for multiple comparisons) and the resulting cluster was used to calculate the displacement error based on the centre of gravity. This process was repeated for 29 areas, which had at least three patients with damage. Finally, the same process was repeated using the lesion principal components (the vascular territories). A Welch *t* test was used to statistically compare the displacement values across the three lesion models.

## RESULTS

### Neuropsychological and lesion distribution

A summary of the patients’ scores is provided in Supplementary Materials 1. The sample contained a range of aphasic performance from global/severe to well-recovered cases. The patients’ lesion overlap map is provided in Figure 1 and primarily covered the left hemisphere area supplied by the MCA (Phan, Donnan, Wright, & Reutens, 2005). There was also evidence of some more general atrophy which may well reflect the effect of long-term vascular load in this patient group. The maximum number of participants who had a lesion in any one voxel was 56 (MNI co-ordinates −38, −9, 24; anatomy of peak: anterior segment of arcuate fasciculus).

### Behaviour factors

The factor analysis on the behavioural data revealed four orthogonal dimensions (see Table 1), which replicates findings from previous studies that had less than half the number of participants (Butler et al., 2014; Halai et al., 2017). The degree to which each test loaded on the factors allows us to interpret the cognitive meaning of each factor. The first factor loaded highest with repetition but also with naming and digit span (and weakly with spoken sentence comprehension) suggesting this factor relates to phonological ability. The second factor loaded highest with picture matching but also with synonym judgement, Camel and Cactus Test and type/token ratio suggesting this factor relates semantic processing. The third factor loaded highest with minimal pairs, Ravens Coloured Progressive Matrices and the Brixton spatial anticipation test suggesting that this factor relates to executive-related problem-solving or decision-making processes. The fourth factor loaded highest with the number of speech tokens produced but also with words per minute and mean length of utterances, which suggests that this factor relates to the quantity of speech produced.

### Lesion principal components

Before reporting the results, it should be noted that the sign (positive or negative) of the coefficients is arbitrary, since variance does not depend on sign, however, the order of the components is important, with the first component accounting for the most variance and the last component the least. The minimum description length criteria (Akaike, 1974; Rissanen, 1983) suggested that there were 20 components within the current dataset. We also used the Kolmogorov information criterion (KIC) which generated an estimate of 32 components within the dataset. For clarity, we only show the results for the 20 component model in the main manuscript but have provided the results of the 32 component model in Supplementary materials. In brief, the cluster pattern remained the same but several big clusters breakdown into smaller, more scattered clusters which are much less interpretable than the 20 component solution (see Supplementary Figure 2 for all 32 components). Minimal description length (MDL) is frequently used in ICA studies for estimation of dimensions and has also been used with PCA. As noted above, this method suggested 20 components for our patient group. We performed a follow up permutation analysis to determine how many components would be estimated by randomly selecting 5, 10, …, 65 and 70 patients from the whole patient group. The permutation test was repeated 100 times for each sample size. The result is provided in Supplementary Figure 3. This analysis showed that when using smaller sample sizes, the number of estimated components vary widely but the number stabilises for larger patient samples. The median component number for a sample size of N=40 was 17, while for N=60 it was 19. For a sample size N=65 and 70, the median value was 20 components. This result suggests that there are likely to be around 20 components and, accordingly, this is the value we used in the study.

**Figure 2.**
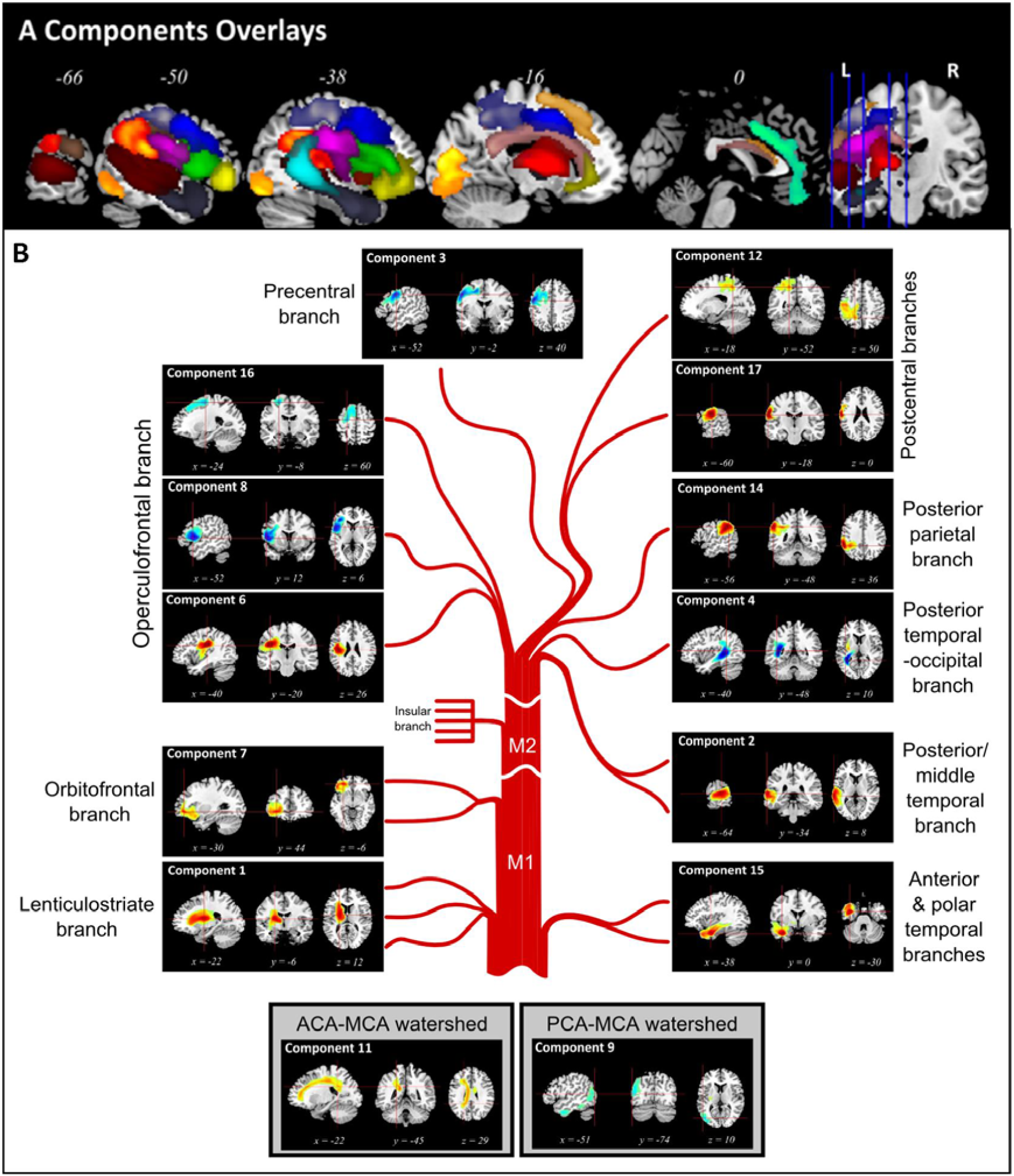
Principal component analysis of abnormality likelihood maps. A: Components overlays. Components 1 – 8, 11 - 17 are shown on the same brain template. B: individual components of MCA illustrated in detail – note the brief labels refer to visually-matched MCA components but may well also reflect other important vascular individual differences such as alternative branching, variable watershed regions, etc. (see main text for descriptions).

**Figure 3.**
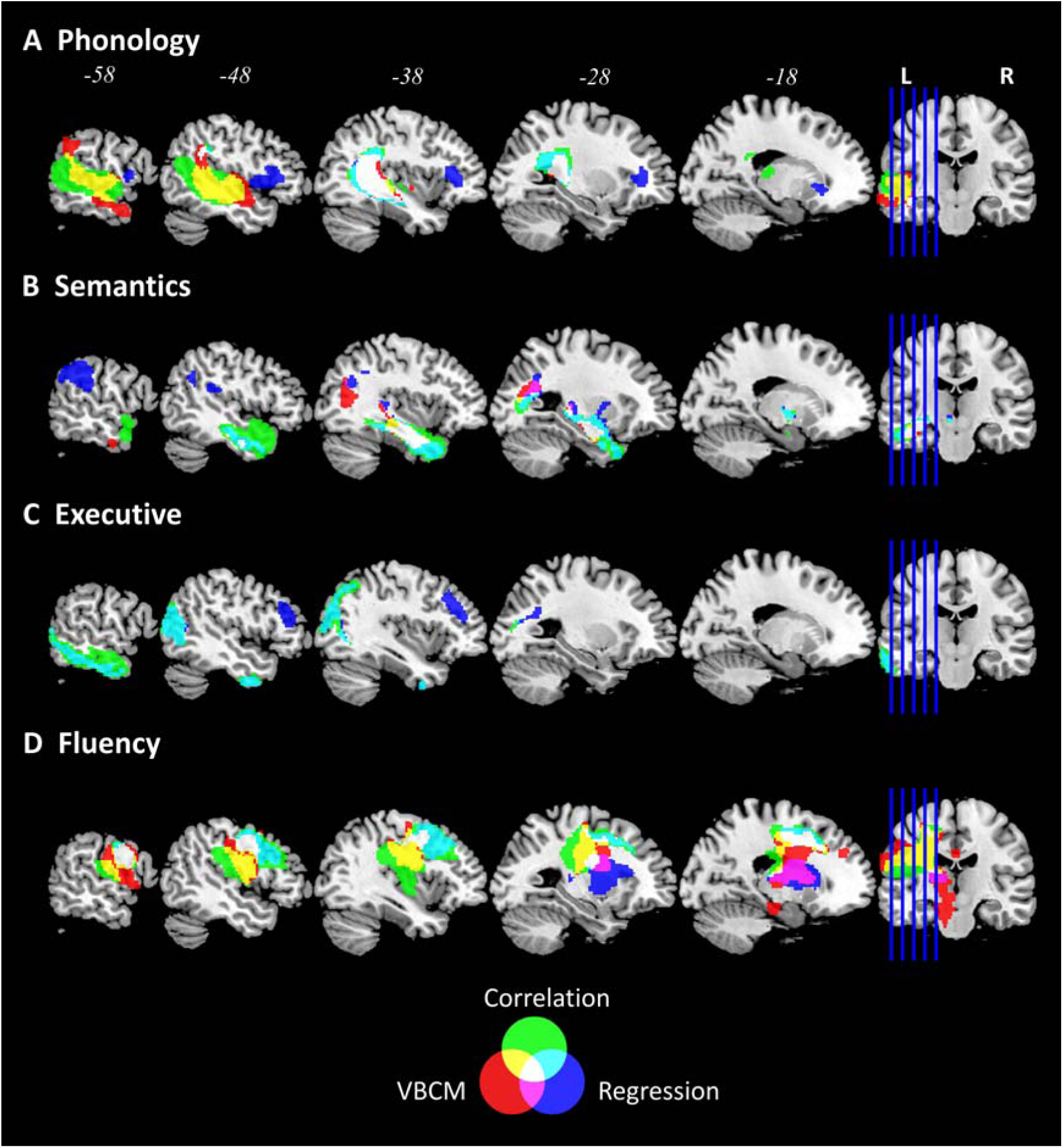
lesion – behaviour mapping. Red colour denotes results of the voxel based correlational analysis with behaviour components. Blue colour denotes regression analysis using lesion components to predict behaviour factor scores. Green colour denotes the Pearson correlation between lesion components and behaviour components. The overlapping regions between the analyses are coloured coded according to the legend (Venn diagram).

The components could be grouped based on location within the fronto-parietal lobes or temporal lobe. Most components belonged to the MCA territory and were located in the fronto-parietal lobes, where Component 1 was subcortical, Components 3, 6, 7, 8, 12, 16, and 17 were frontal and Component 14 was parietal. There were three components in the temporal lobe (Components 2, 4, and 15). Two or three components were located along the watershed between territories (Components 9 and 11, and possibly 16). Component 13 fell within the medial frontal region and was likely to belong to the anterior cerebral artery territory and Component 9 in the occipital lobe was likely to belong the posterior cerebral artery territory. The four other components were either scattered or included regions in the right hemisphere and thus hard to interpret (Components 10, 18, 19, 20. See Supplementary Figure 1 for all 20 components). A visual illustration of the components overlay is shown in Figure 2 which arranged the lesion components based on the vasculature (a detailed description components’ locations were given in Supplementary Table 2. In the following paragraphs, we describe the location of each cluster in relation to anatomy using the Harvard-Oxford (grey matter) and natbrainlab (white matter) atlases. Given that the seminal angiography studies of the MCA sub-branches were conducted before 3D brain mapping had been developed we visually compared the PCA-derived clusters and the hand-drawn maps from Michotey et al (1974). It is important to note that the PCA will also detect other sources of consistent individual differences in vascular supply and damage that is related to individual variations in vascular branching, patterns of vascular collateralisation, and so on. Perhaps most importantly for the study it is worth noting that the key aim of this approach is to investigate the possibility of moving away from voxel-based analysis to one that embraces the collinearities in voxel damage. In this context it is more powerful that the PCA outcome is a reflection of all sources of variance found in the patient sample in addition to the typical vascular branching itself.

The first component was located in subcortical territories that mainly occupy the putamen and thalamus, and underlying white matter, e.g. the cortical spinal and the inferior occipito-frontal fasciculus. Thus, it corresponded to the lenticulostriate branch, originating from the M1 segment of the MCA. The current data set did not discover a pure insular branch, which originates from the M2 segment of the MCA. Components 8 and 6 corresponded to the operculofrontal branch, originating from the M3 segment. Component 8 was located at the frontal operculum cortex, extending to inferior frontal gyrus (pars opercularis) and precentral gryus. Component 6 was located at the central opercular cortex, extending to insular and parietal operculum cortex. Component 7, 3, 17, and 12 corresponded to the orbitofrontal, precentral, central, and postcentral branches, respectively. These four are the territories of the frontal cortical branches, originating from the M4 segment. Component 7 came after the subcortical branch, located at the orbital frontal cortex, extending to frontal pole and inferior frontal gyrus (pars triangularis). Following it was Component 3, located at the middle frontal gyrus, extending to the precentral gyrus and inferior frontal gyrus. Component 17 was lateral to Component 3, lying on the central sulcus, occupying both precentral and postcentral gyri. Component 12 was posterior to Component 3 and was located at the postcentral gyrus, but occupying a large section of the superior parietal lobule. Component 16 was located at the middle and superior frontal gryus, and was superior to the other frontal components. It is possible that this either reflected the most dorsal frontal branch, or belonged to the watershed with the anterior cerebral artery. Component 14 was completely within the parietal lobe, including supramarginal gyrus and angular gyrus and was linked to the posterior parietal branch.

Moving inferiorly into the temporal lobe territories, we found Component 2 which was in the lateral temporal lobe, extending from the anterior superior and middle temporal gyrus to the angular gyrus and supramarginal gyrus. As the peak was posterior, we suggest that this reflects the posterior temporal branch. The medial temporal lobe structures made up Component 4, including the inferior longitudinal fasciculus and posterior segment of the arcuate fasciculus, therefore was likely to reflect the posterior temporal-occipital branch. The anterior inferior temporal lobe made up Component 15, including the anterior portion of the inferior longitudinal fasciculus. The one component in the occipital lobe (Component 5) was located over the inferior division of lateral occipital cortex, occipital fusiform gyrus, and occipital pole, and therefore was more likely to belong to the posterior cerebral artery.

We labelled Components 9 and 11 as watershed components as they overlapped with the territory between cerebral arteries (Michotey et al., 1974). Component 9 overlapped with the watershed territory between MCA and posterior cerebral artery, occupying the lateral occipital cortex down to the middle and inferior temporal gyrus. Although both Component 5 and 9 are located within occipital regions we do not believe they reflect similar underlying territories as Component 5 is more focal compared to Component 9. Component 11 overlapped with the watershed territory between MCA and anterior cerebral artery, occupying white matters including the corpus callosum and cingulum. This component also extends into the right ventricle and therefore might also capture atrophy related to the effects of long-standing vascular load. Component 10 may partially reflect the cortical portion of the watershed between anterior cerebral artery and MCA, as it covers a range of regions along the frontal pole and middle frontal gyrus but the cluster in the right hemisphere makes it difficult to interpret.

### Displacement error using different lesion models

We simulated the ‘ground truth’ for three different lesion models (single voxel, Brodmann areas and vascular territories) and found that the mean displacement error was largest for the Brodmann atlas (22.46 mm, SD = 16.28 mm). This was followed by the single voxel model (16.34 mm, SD = 6.87 mm) and then the vascular territories model (13.85 mm, SD = 4.65 mm). The displacement of the vascular territories was significantly smaller than the single voxel simulation (Welch’s *t* test, *t* = 2.14, *p* = 0.0493, two-tailed) and the Brodmann atlas simulation (Welch’s *t* test, *t* = 2.66, *p* = 0.0117, two-tailed). See Supplementary Figure 4 for a violin plot of the displacement error for each model and a 3D brain projected figure showing which areas of the have high/low displacement (for reference, we also show the probability of damage following an MCA stroke by Phan et al., 2005).

### Voxel-wise neural correlates of the behaviour factors

The neural correlates identified using VBCM for phonology, semantics and fluency are shown in Figure 3 (red regions) and described in Supplementary Table 3 (Alphasim corrected *p* < 0.01 with voxel *p* < 0.001). There were no significant clusters for the executive factor at this statsitical threshold. The cluster related to phonology was found in the dorsal auditory language pathway, including middle and superior temporal gyrus, extending posterior to supramarginal gyrus and medial whiter matter substrates, e.g., inferior longitudinal fasciculus and arcuate posterior segment (5641 voxels; peak coordinates: −64, −18, −12; peak *r* = 0.55, *p* < 0.001). Semantic ability was related to two clusters. The first one covered the lateral and medial anterior temporal lobe and extending to medial temporal lobe inferior longitudinal fasciculus (1175 voxels; peak coordinates: −36, −12, −18; peak *r* = 0.54, *p* < 0.001). The second cluster was at the posterior end of the inferior longitudinal fasciculus and extended into the superior division of lateral occipital cortex (509 voxels; peak coordinates: −34, −66, 18; peak *r* = 0.46, *p* < 0.001). Finally, the fluency factor correlated with a large cluster in the frontal lobe, covering the precentral and postcentral gyrus, thalamus, central opercular cortex, and also white matter including the frontal aslant tract, cortico spinal, corpus callosum, and cingulum (9479 voxels; peak coordinates: −60, −6, 20; peak *r* = 0.57, *p* < 0.001).

### Relating lesion component scores to the behavioural factor scores

The Pearson correlation analysis (Figure 3 green regions) between lesion components and behaviour components showed that the correlated components overlapped with the VBCM results. In detail, Component 2 (*r* = −0.409) and 4 (*r* = 0.419) correlated with phonology components score, Component 15 (*r* = −0.495) correlated with semantics, Component 9 (*r* = 0.477) correlated with executive and lastly, Component 3 (*r* = 0.436) and 6 (*r* = −0.407) correlated with fluency (all *p* < 0.0006).

We used stepwise regression to determine if the lesion territory components can be used to predict behavioural factors. The results are shown in Figure 3 (blue clusters) and subsequent details for the clusters in Supplementary Table 4. We found a significant model for predicting phonological ability (*F*(2, 67) = 10.288, *p* < 0.001), which included Component 4 (beta = 0.429) and Component 8 (beta = 0.244), with an adjusted *r* square of 0.212. We mapped the *z*-transformed coefficients back into brain space and found two significant clusters. The largest cluster was located over the left inferior longitudinal fasciculus and the posterior segment of the arcuate fasciculus, while the second cluster was located in the left pars opercularis and frontal operculum cortex. A significant model predicted semantic ability (*F*(3, 66) = 11.640, *p* < 0.001), which included Component 15 (beta = – 0.418), Component 14 (beta = – 0.336) and Component 12 (beta = 0.256), with an adjusted *r* square of 0.316. We found three significant clusters on the brain, of which the largest cluster was located within the left temporal lobe and the inferior longitudinal fasciculus. The second cluster was identified in the left supramarginal gyrus and the third cluster was in the left optic radiations. We observed a significant model for executive ability (*F*(2, 67) = 14.152, *p* < 0.001), which included Component 9 (beta = 0.443) and Component 10 (beta = −0.266), with an adjusted *r* square of 0.276. Neural regions related to the executive factor were located in the watershed regions. The largest cluster overlapped with inferior temporal lobe running posterior to the occipital lobe, while the second cluster was located within the left middle frontal gyrus, the frontal pole and the inferior frontal gyrus. Finally, a significant model was identified for speech fluency ability (*F*(2, 67) = 11.665, *p* < 0.001), which included Component 3 (beta = 0.352) and Component 1 (beta = – 0.274), with an adjusted *r* square of 0.236. The neural results revealed two significant clusters, the largest of which was located in the left middle frontal gyrus and precentral gyrus. The smaller cluster was found in subcortical regions such as the cortico-spinal tract, putamen and thalamus.

## DISCUSSION

Modern neuroimaging now allows sophisticated *in vivo* lesion mapping. Many previous studies using lesion-symptom mapping have not taken into account the non-random distribution of brain damage after stroke, which is constrained by the vasculature. In the present study, we showed that principal component analysis of the lesions in post-stroke aphasia can reveal the underlying components of damage across the MCA territory, which are striking similarity to the vascular structure and forms of vascular individual variation identified by seminal angiography studies (Michotey et al., 1974). This parcellation produced less displacement during lesion-symptom mapping inference compared to a standard single voxel model and an anatomical atlas, which suggests that taking into account the vascular territories might improve our understanding of the locus of behavioural deficits in stroke patients. In a secondary stage, we undertook a preliminary exploration of how these lesion components might relate to the patients’ language and cognitive variations.

### Comparing the lesion principal components with angiography

It is striking that the majority of components, identified by varimax-rotated PCA of the neural data, were clusters that reflect many of the cortical and subcortical MCA territories (see Figure 2) identified in classical angiography studies as well as other forms of vascular individual differences (Michotey et al., 1974). We should note that we labelled the PCA clusters by visual comparison with the descriptions provided in these previous seminal studies. These classical studies were conducted long before formal methods had been developed for 3D brain mapping that would allow definitive direct comparisons between the vascular supply of the MCA sub-branches and PCA-derived clusters. As expected, the lesion components showed an anterior-posterior separation covering frontal and temporoparietal regions, respectively. Previous angiography studies identified four branches in the frontal lobe (orbitofrontal, prefrontal, precentral, and central) which were reflected in six PCA components across both lateral and medial frontal regions. We found direct correspondences for the two parietal branches (Components 12 and 14). Component 12 covered superior parietal lobule and extends into postcentral gyrus, while Component 14 was situated over the angular and supramarginal gyri. There was also good similarity with respect to the classical, angiographically-determined MCA temporal territories (temporo-occipital, posterior, middle, anterior temporal, and temporal polar). The PCA identified three components along the caudal-rostral axis (Components 2, 4 and 15). One possible reason for detecting larger temporal principal components in the lesion maps than expected by the angiography studies may be due to the fact that our patient distribution is such that we have relatively fewer cases with damage to the temporal lobe than frontal lobe. This, in turn, may reflect the fact that the middle and ventral parts of the temporal lobe are less likely to be damaged following an MCA infarct: Phan and colleagues (2005) identified that the likelihood of damage dropped to around 10-15% in the middle temporal gyrus and lower than 5% from the inferior temporal gyrus and below. In contrast, the probability of frontal-insular damage is greater than 30%. It is also possible that the hierarchical branching and diameter of the sub-branches is such that certain combinations are likely to be jointly occluded (e.g., the polar and anterior temporal branches). The lesion PCA also identified two other major features of the vascular supply. First, as well as the MCA cortical territories, we also obtained the lenticulostriate branch (Component 1) as well as the watershed territories between MCA-PCA (Component 9) and MCA-ACA (Component 11) which are known to show individual variability (Michotey et al., 1974) and thus these regions may be less likely to emerge in lesion-symptom mapping analyses. Secondly, the analyses also identify more widespread atrophic changes which may well reflect the effects of long-term vascular load.

We note that, inescapably, the spatial maps of principal components depend on the number of the components extracted. When a different optimization criterion was applied, a larger estimated number of components was derived yet the maps relating to these components were much harder to interpret and did not relate obviously to the known cortical vascular supplies (see Supplementary Figure 2). The number of components was also much lower than the number of regions found in many brain parcellation approaches based on cytoarchitecture, function or connectivity. We suspect that this variation reflects differences in the inherent granularity of the raw data in each case. Our estimate of around 20 vascular components aligns with the known number of cortical vascular branches of the MCA (N=12) plus individual differences in vascular supply. In contrast, there are numerous variations in cytoarchitecture or cortical functions, and thus it seems inevitable that there will be many more regions in those resultant parcellations of the brain. Future studies that have access to much larger lesion databases can assess that the spatial clusters identified in this study are efficient in capturing the post-stroke damage, by evaluating predictive modelling of out-of-sample data.

### Impact of vascular territory parcellations in lesion-symptom mapping

Our results showed that the vascular territories parcellation produced smaller displacement to the simulated ground truth than a single voxel or Brodmann map model. The displacement values for the single voxel model and Brodmann areas derived in the current study are very similar to recent studies (i.e. Mah et al., 2014; Sperber & Karnath, 2018). By projecting the displacement values to the brain, we identified that the regions with greatest displacement were typically located on the edge of the MCA territory between the other two major arteries (Michotey et al., 1974). We offer two possible explanations for the increased displacement in these regions: 1) the number of lesions in these territories is lower and thus the estimations are noisier; or 2) the boundaries between watershed territories are inherently variable across individuals, especially as dual supplied regions (such as the anterior temporal lobe and orbito-frontal cortex) would be more robust. In Supplementary Figure 4C, we illustrate that the most displaced areas are in the regions that are less likely to be damaged in an MCA stroke population (Phan et al., 2005). This suggests that we may need to either include a very large population of patients to capture the <10% probability of damage to these regions sufficiently and/or take into cases with strokes related to the other major arteries in conjunction with the MCA.

### Comparing the neural correlates of behaviour at the voxel and lesion territory level

By way of a preliminary exploration, we compared voxel-based analyses (VBCM) with the results from the lesion component to behavioural mapping. The results for phonological ability showed overlap between both methods in the posterior temporal region including the underlying posterior segment of the arcuate fasciculus, which has been shown to be involved in the dorsal auditory language pathway (Catani & Jones, 2005; Hickok & Poeppel, 2004, 2007; Parker et al., 2005; Saur et al., 2008; Ueno, Saito, Rogers, & Lambon Ralph, 2011). This region has also been implicated using other methodologies, e.g. diffusion-weighted imaging and tractography (Glasser & Rilling, 2008; Park et al., 2011), intraoperative subcortical electrical stimulation (Duffau, Gatignol, Mandonnet, Capelle, & Taillandier, 2008; Leclercq et al., 2010), neuroanatomically-constrained computational models (Ueno et al., 2011), and VLSM analysis (Bates et al., 2003; Fridriksson, Richardson, Baker, & Rorden, 2011). There were some differences between the two methods. The VBCM analysis revealed correlations with a larger region including the lateral superior temporal gyrus and middle temporal gyrus, which is associated with the ventral language pathway (Hickok & Poeppel, 2004, 2007; Price, 2010). Although there is a lesion component (no.4) that overlaps with this extended area, it did not appear in the regression analysis as the posterior temporal component (no. 2) explained a greater amount of variance and thus, Component 4 did not enter the regression analysis. Consistent with this conclusion, the simple correlation analysis identified a larger temporal lobe region as correlated with phonological ability (Component 2 in addition to Component 4) which overlapped considerably with the area from the voxel-based analysis (see yellow-marked area in Figure 3A). The lesion-component regression analysis also detected a second critical region for phonological abilities within the inferior frontal region. This follows from the fact that, having accounted for the variance along the temporal lobe, the regression analysis was able to identify additional independent variance associated with the inferior frontal component, which was not found in the VBCM or the simple correlations. Indeed, the inferior frontal lobe has been repeatedly linked with phonological processing in previous studies, and has been associated with mapping sound structure to production and/or executive-attention mechanisms in selecting the correct sound mapping (Price, 2010; Rauschecker & Scott, 2009; Vigneau et al., 2006).

VBCM and the behaviour-lesion components regression both revealed clusters in the white matter of anterior temporal lobe when mapping semantic ability. This finding converges with evidence that the anterior temporal lobe is important for successful semantic representation (Lambon Ralph, Jefferies, Patterson, & Rogers, 2016). The anterior temporal lobe has been suggested to act as a transmodal semantic hub, as revealed by multiple convergent methodologies, e.g., semantic dementia (Acosta-Cabronero et al., 2011; Patterson, Nestor, & Rogers, 2007), repetitive transcranial magnetic stimulation (Pobric, Jefferies, & Lambon Ralph, 2007), fMRI (Coutanche & Thompson-Schill, 2014; Peelen & Caramazza, 2012; Visser, Jefferies, & Lambon Ralph, 2010), and resting-state fMRI (Zhao et al., 2017). Studies investigating the white matter connectivity, associated with the anterior temporal lobe have revealed that the uncinate fasciculus, inferior longitudinal fasciculus and inferior frontooccipital fasciculus are related to semantic performance (Acosta-Cabronero et al., 2011; Han et al., 2013; Mandonnet, Nouet, Gatignol, Capelle, & Duffau, 2007). The lesion structure regression analysis was able to detect an extra region located within supramarginal and angular gyrus. These ventral parietal regions have be suggested to be related to semantic processing (Binder, Desai, Graves, & Conant, 2009; Hartwigsen, Golombek, & Obleser, 2015). However, there is growing recent evidence that the role of the angular gyrus might not be selective for semantic processing (Humphreys, Hoffman, Visser, Binney, & Lambon Ralph, 2015), but rather may serve numerous cognitive domains (Humphreys & Lambon Ralph, 2014), including episodic memory (Humphreys & Lambon Ralph, 2014; Rugg & Vilberg, 2013; Wagner, Shannon, Kahn, & Buckner, 2005).

Speech fluency is usually the first domain on which aphasia patients are categorised (Goodglass & Kaplan, 1983). In our previous behavioural work, we found that the quantity of speech produced can be separated from phonological and semantic abilities (Halai et al., 2017). Classically, the inferior frontal gyrus (termed Broca’s area) had been associated with poor speech production (Broca, 1861); however, recent studies have areas within the insula, precentral gyrus and subcortical structures such as the caudate, putamen and thalamus have been implicated in speech fluency (Basilakos et al., 2014; Bates et al., 2003; Blank, Scott, Murphy, Warburton, & Wise, 2002; Halai et al., 2017). The current study found overlap between the two lesion-behaviour mapping methodologies, both identifying the precentral gyrus, middle frontal gyrus and thalamus. VBCM also found a wider region over the frontal lobe that extended laterally into inferior frontal gyrus as well as medially down the cortico-spinal tract. As with phonological ability, the broader region identified by VBCM was also found in the simple correlations (see yellow-marked region in Fig.3D) – again reflecting the fact that in stepwise regression only the components that capture the unique sources of variance are included, whilst any components that have similar but slightly weaker explanatory power are disregarded.

Finally, VBCM analysis failed to identify any neural correlates for the executive factor (as also found in previous studies, unless low statistical thresholds are applied (Butler et al., 2014; Halai et al., 2017; Lacey et al., 2017). In contrast, the lesion component regression analysis identified regions at the edge of the MCA territory that predicted executive abilities. These identified regions align closely with the proposed frontoparietal multi-demand network for executive function (e.g. Duncan, 2010). Furthermore, studies that extract network measures from resting state fMRI and DTI have found a large scale network of regions across the frontal, parietal and temporal lobe related to executive functions (Li et al., 2009; van den Heuvel, Stam, Kahn, & Pol, 2009). This distributed nature converges with the results observed in our analysis, whereby areas across all lobes are found to be predictive of better executive performance.

## Conclusion

Studies of cognitive and language impairments in stroke patients have provided valuable insights into understanding the organisation of the brain and provide implications for clinical application. In the present study, we have demonstrated that the underlying structure of the lesions reflects the brain’s vasculature that can be utilised in lesion-symptom mapping to reduce mislocation error in stroke patients. Future, formal studies will be able to test and validate the use of these PCA-derived lesion components in lesion-symptom mapping, prediction modelling and related neuropsychological approaches that aim to relate brain to behaviour.

## Acknowledgements

We thank all the patients and carers for their continued support of our research programme. This study was supported by an ERC advanced grant (GAP: 670428 - BRAIN2MIND_NEUROCOMP) and MRC programme grants (MR/R023883/1; MR/R023883/1) to MALR, and from the Rosetrees Trust (A1699) to ADH and MALR.

