## Supplementary figures and images for "Evaluating the granularity and statistical structure of lesions and behaviour in post-stroke aphasia"

### Supplementary Figure 1

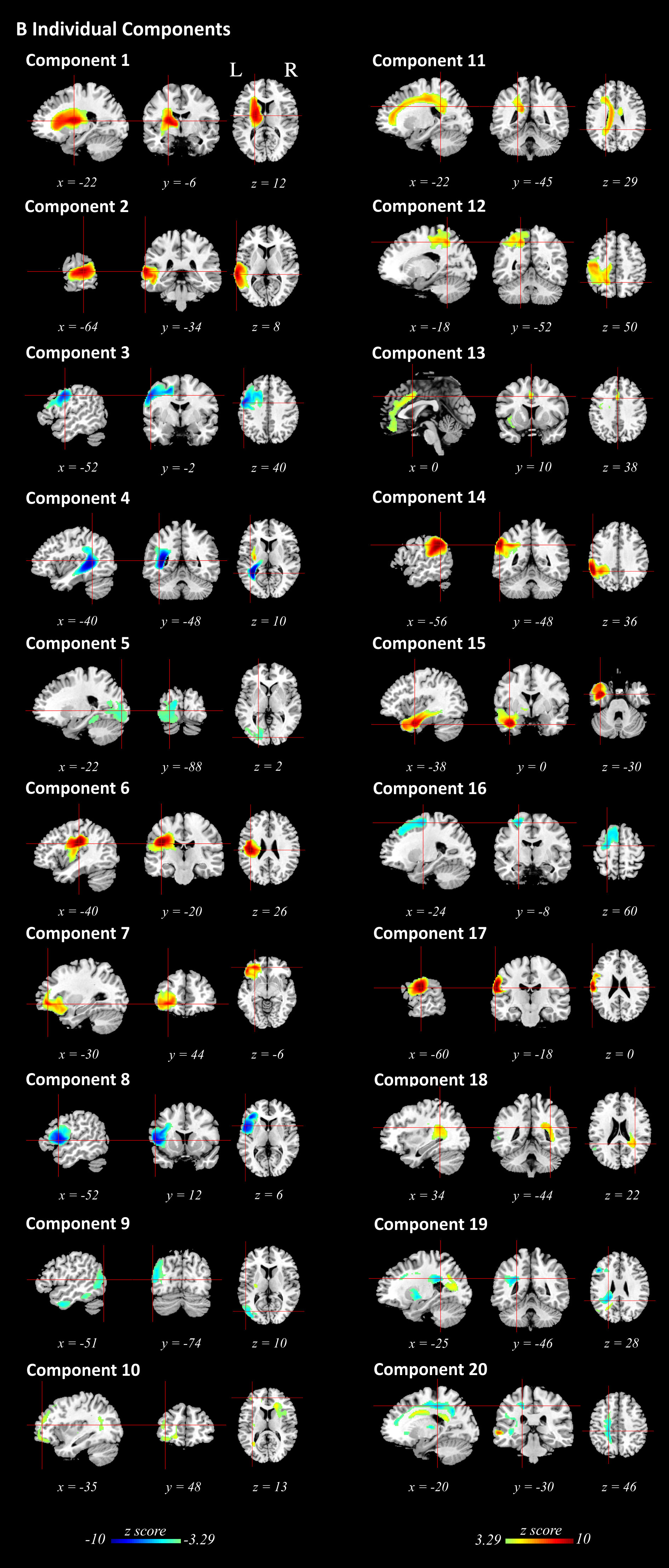

### Supplementary Figure 2

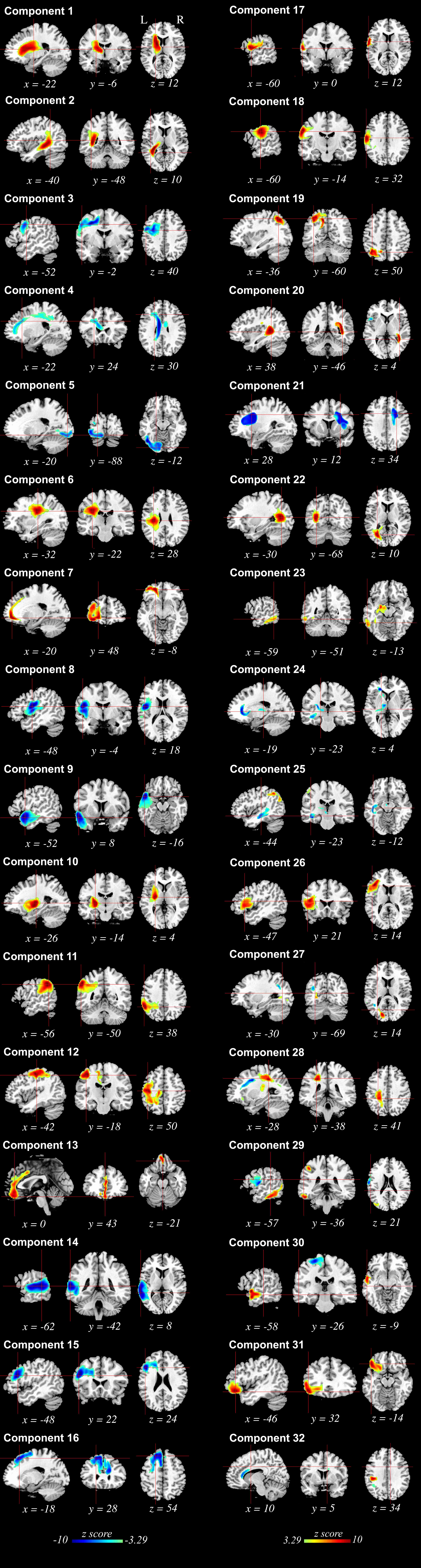

### Supplementary Figure 3

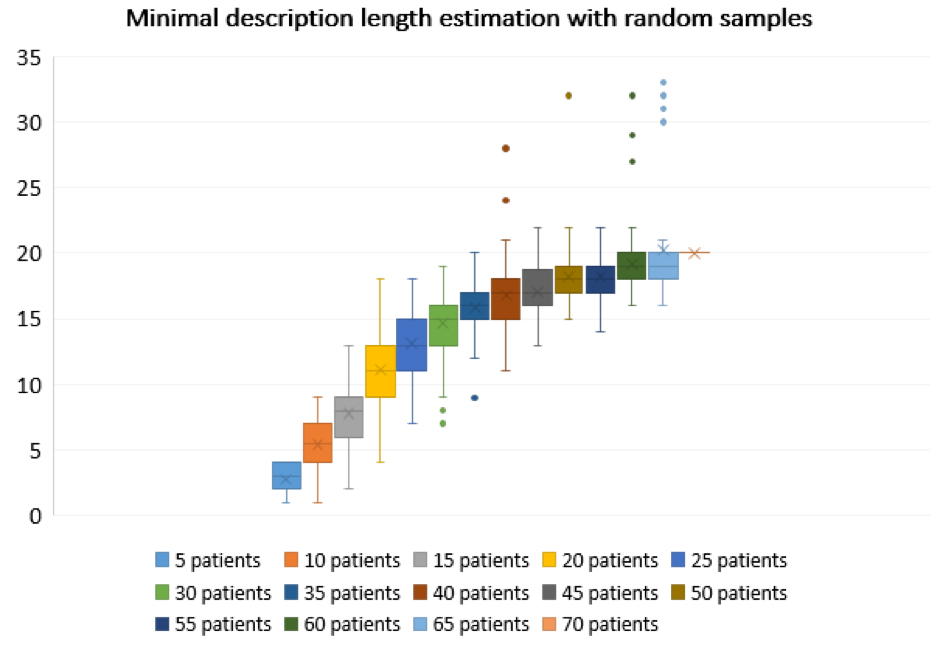

### Supplementary Figure 4

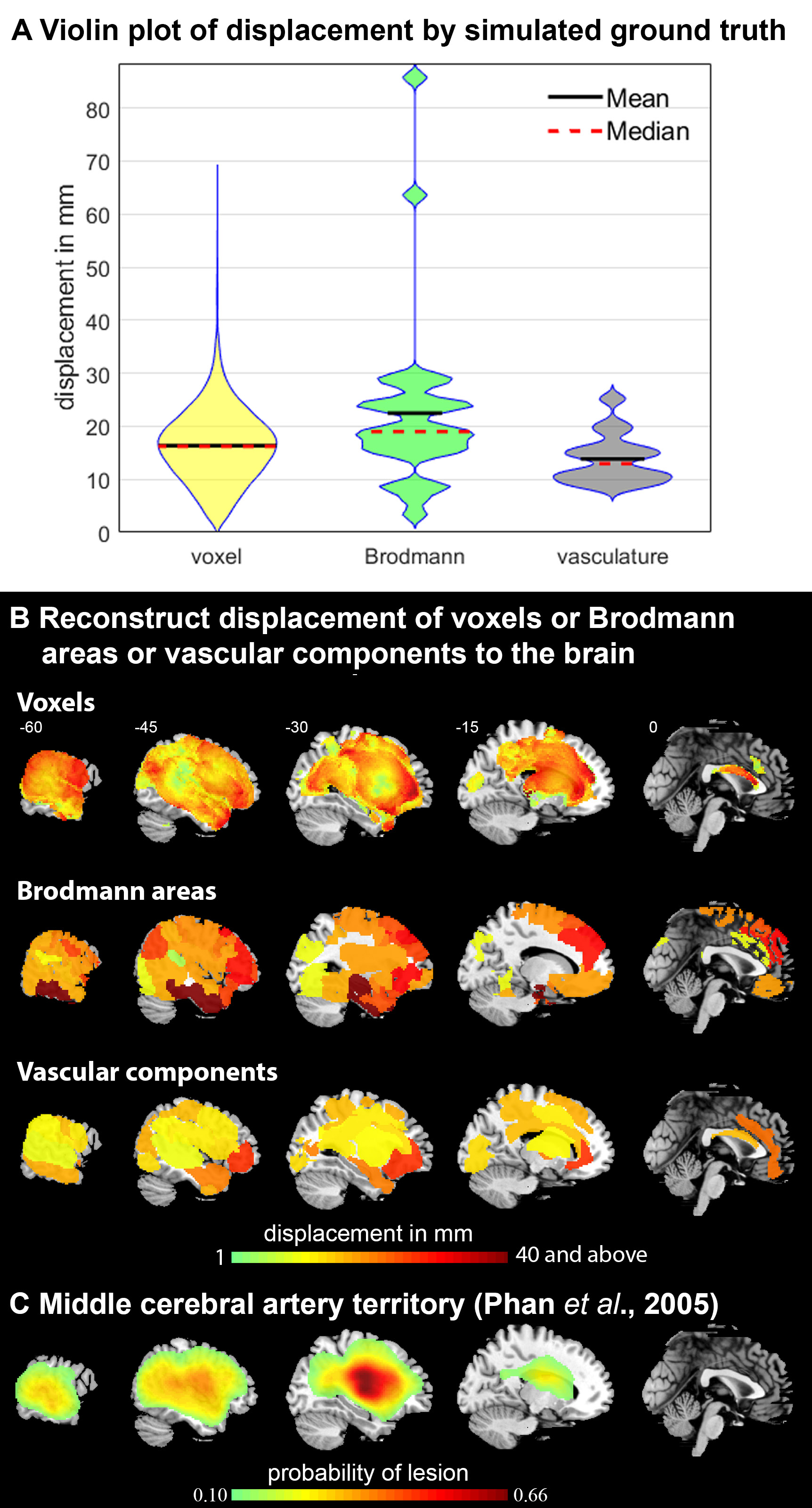
