## Supplementary Table 2 for "Evaluating the granularity and statistical structure of lesions and behaviour in post-stroke aphasia"

**Table 2. Lesion components location**

| PCA  Components | | Size  (voxels) | Percent  On template | Peak Coordinates  (MINI x, y, z) | Peak  (*z*) |
| --- | --- | --- | --- | --- | --- |
| **Component 1** | | **5191** |  | **-22 -6 12** | **9.20** |
| G | Putamen (L) | 685 | 74% |  |  |
|  | Thalamus (L) | 494 | 36% |  |  |
|  | Caudate (L) | 373 | 66% |  |  |
|  | Insular Cortex (L) | 252 | 22% |  |  |
|  | Pallidum (L) | 186 | 60% |  |  |
| W | Cortico Spinal (L) | 1178 | 33% |  |  |
|  | Inferior Occipito Frontal Fasciculus (L) | 455 | 33% |  |  |
|  | Corpus Callosum (L) | 438 | 8% |  |  |
|  | Internal Capsule (L) | 322 | 26% |  |  |
|  | Fornix (L) | 284 | 23% |  |  |
|  | Uncinate (L) | 198 | 22% |  |  |
| **Component 2** | | **4916** |  | **-64 -34 8** | **9.67** |
| G | Superior Temporal Gyrus, posterior division (L) | 814 | 93% |  |  |
|  | Middle Temporal Gyrus, temporooccipital part (L) | 560 | 66% |  |  |
|  | Middle Temporal Gyrus, posterior division (L) | 505 | 40% |  |  |
|  | Planum Temporale (L) | 405 | 76% |  |  |
|  | Angular Gyrus (L) | 290 | 31% |  |  |
|  | Supramarginal Gyrus, posterior division (L) | 285 | 27% |  |  |
|  | Heschls Gyrus (includes H1 and H2) (L) | 208 | 65% |  |  |
|  | Lateral Occipital Cortex, superior division (L) | 151 | 3% |  |  |
|  | Planum Polare (L) | 107 | 30% |  |  |
|  | Superior Temporal Gyrus, anterior division (L) | 102 | 40% |  |  |
| W | Arcuate Posterior Segment (L) | 243 | 33% |  |  |
|  | Inferior Longitudinal Fasciculus (L) | 208 | 12% |  |  |
| **Component 3** | | **4946** |  | **-52 -2 40** | **-8.16** |
| G | Middle Frontal Gyrus (L) | 1519 | 55% |  |  |
|  | Precentral Gyrus (L) | 1367 | 32% |  |  |
|  | Inferior Frontal Gyrus, pars opercularis (L) | 180 | 25% |  |  |
| W | Corpus Callosum (L) | 240 | 4% |  |  |
|  | Cortico Spinal (L) | 229 | 7% |  |  |
| **Component 4 Cluster 1** | | **3043** |  | **-34 -48 10** | **-9.98** |
| W | Inferior Longitudinal Fasciculus (L) | 610 | 35% |  |  |
|  | Arcuate Posterior Segment (L) | 398 | 53% |  |  |
|  | Cortico Spinal (L) | 359 | 10% |  |  |
|  | Optic Radiations (L) | 310 | 57% |  |  |
|  | Corpus Callosum (L) | 201 | 3% |  |  |
|  | Fornix (L) | 151 | 12% |  |  |
|  | Inferior Occipito Frontal Fasciculus (L) | 147 | 11% |  |  |
|  | Internal Capsule (L) | 139 | 11% |  |  |
| **Component 4 Cluster 2** | | **1091** |  | **-30 -16 6** | **7.01** |
| G | Putamen (L) | 420 | 46% |  |  |
|  | Pallidum (L) | 147 | 47% |  |  |
|  | Insular Cortex (L) | 123 | 11% |  |  |
| W | Inferior Occipito Frontal Fasciculus (L) | 128 | 9% |  |  |
| **Component 5** | | **3107** |  | **-14 -90 10** | **-5.10** |
| G | Lateral Occipital Cortex, inferior division (L) | 788 | 39% |  |  |
|  | Occipital Fusiform Gyrus (L) | 387 | 42% |  |  |
|  | Occipital Pole (L) | 374 | 14% |  |  |
|  | Lingual Gyrus (L) | 253 | 17% |  |  |
|  | Inferior Temporal Gyrus, temporooccipital part (L) | 124 | 18% |  |  |
| W | Inferior Occipito Frontal Fasciculus (L) | 230 | 17% |  |  |
|  | Corpus Callosum (L) | 206 | 4% |  |  |
| **Component 6** | | **4315** |  | **-40 -20 26** | **10.39** |
| G | Central Opercular Cortex (L) | 681 | 69% |  |  |
|  | Postcentral Gyrus (L) | 334 | 9% |  |  |
|  | Insular Cortex (L) | 247 | 21% |  |  |
|  | Parietal Operculum Cortex (L) | 129 | 22% |  |  |
|  | Precentral Gyrus (L) | 122 | 3% |  |  |
| W | Cortico Spinal (L) | 710 | 20% |  |  |
|  | Arcuate Anterior Segment (L) | 482 | 90% |  |  |
|  | Long Segment (L) | 255 | 72% |  |  |
|  | Corpus Callosum (L) | 121 | 2% |  |  |
| **Component 7** | | **3900** |  | **-30 44 -6** | **8.18** |
| G | Frontal Pole (L) | 1129 | 16% |  |  |
|  | Frontal Orbital Cortex (L) | 1078 | 63% |  |  |
|  | Inferior Frontal Gyrus, pars triangularis (L) | 225 | 37% |  |  |
|  | Insular Cortex (L) | 203 | 18% |  |  |
| W | Corpus Callosum (L) | 213 | 4% |  |  |
|  | Uncinate (L) | 128 | 14% |  |  |
| **Component 8** | | **4718** |  | **-52 12 6** | **-8.71** |
| G | Precentral Gyrus (L) | 649 | 15% |  |  |
|  | Inferior Frontal Gyrus, pars opercularis (L) | 590 | 81% |  |  |
|  | Central Opercular Cortex (L) | 418 | 42% |  |  |
|  | Frontal Operculum Cortex (L) | 347 | 96% |  |  |
|  | Insular Cortex (L) | 316 | 27% |  |  |
|  | Inferior Frontal Gyrus, pars triangularis (L) | 249 | 41% |  |  |
| W | Inferior Occipito Frontal Fasciculus (L) | 137 | 10% |  |  |
|  | Arcuate Anterior Segment (L) | 115 | 21% |  |  |
| **Component 9 Cluster 1** | | **3455** |  | **-66 -24 -14** | **-6.06** |
| G | Lateral Occipital Cortex, superior division (L) | 923 | 19% |  |  |
|  | Middle Temporal Gyrus, posterior division (L) | 588 | 46% |  |  |
|  | Lateral Occipital Cortex, inferior division (L) | 514 | 25% |  |  |
|  | Inferior Temporal Gyrus, posterior division (L) | 318 | 32% |  |  |
|  | Middle Temporal Gyrus, anterior division (L) | 250 | 56% |  |  |
|  | Middle Temporal Gyrus, temporooccipital part (L) | 235 | 28% |  |  |
|  | Inferior Temporal Gyrus, anterior division (L) | 204 | 62% |  |  |
|  | Inferior Temporal Gyrus, temporooccipital part (L) | 135 | 19% |  |  |
|  | Temporal Pole (L) | 113 | 5% |  |  |
| **Component 9 Cluster 2** | | **597** |  | **-44 34 24** | **-5.66** |
| G | Middle Frontal Gyrus (L) | 440 | 16% |  |  |
| **Component 10 Cluster 1** | | **2072** |  | **16 20 22** | **5.08** |
| W | Corpus Callosum (R) | 442 | 8% |  |  |
|  | Internal Capsule (R) | 299 | 13% |  |  |
| **Component 10 Cluster 2** | | **1637** |  | **-16 44 -10** | **4.99** |
| G | Frontal Pole (L) | 473 | 7% |  |  |
|  | Putamen (L) | 200 | 22% |  |  |
|  | Middle Frontal Gyrus (L) | 153 | 6% |  |  |
|  | Frontal Orbital Cortex (L) | 135 | 8% |  |  |
|  | Pallidum (L) | 125 | 40% |  |  |
| W | Anterior Commissure (L) | 163 | 27% |  |  |
|  | Cortico Spinal (L) | 110 | 3% |  |  |
|  | Uncinate (L) | 106 | 12% |  |  |
| **Component 11** | | **5598** |  | **-16 -46 22** | **6.84** |
| W | Corpus Callosum (L) | 1589 | 28% |  |  |
|  | Cingulum (L) | 1187 | 24% |  |  |
|  | Cortico Spinal (L) | 611 | 17% |  |  |
|  | Corpus Callosum (R) | 222 | 4% |  |  |
|  | Cingulum (R) | 100 | 2% |  |  |
| **Component 12** | | **5530** |  | **-18 -52 50** | **7.33** |
| G | Postcentral Gyrus (L) | 1556 | 43% |  |  |
|  | Superior Parietal Lobule (L) | 904 | 62% |  |  |
|  | Precentral Gyrus (L) | 635 | 15% |  |  |
|  | Precuneous Cortex (L) | 135 | 5% |  |  |
|  | Supramarginal Gyrus, posterior division (L) | 108 | 10% |  |  |
| W | Cortico Spinal (L) | 301 | 9% |  |  |
|  | Corpus Callosum (L) | 275 | 5% |  |  |
|  | Cingulum (L) | 212 | 4% |  |  |
| **Component 13** | | **999** |  | **-2 10 38** | **5.54** |
| G | Paracingulate Gyrus (R) | 221 | 16% |  |  |
|  | Cingulate Gyrus, anterior division (R) | 183 | 14% |  |  |
|  | Cingulate Gyrus, anterior division (L) | 104 | 10% |  |  |
| **Component 14** | | **4629** |  | **-56 -48 36** | **9.98** |
| G | Supramarginal Gyrus, anterior division (L) | 685 | 72% |  |  |
|  | Supramarginal Gyrus, posterior division (L) | 671 | 64% |  |  |
|  | Angular Gyrus (L) | 548 | 58% |  |  |
|  | Parietal Operculum Cortex (L) | 495 | 86% |  |  |
|  | Planum Temporale (L) | 229 | 43% |  |  |
|  | Lateral Occipital Cortex, superior division (L) | 131 | 3% |  |  |
| W | Arcuate Posterior Segment (L) | 441 | 59% |  |  |
|  | Cortico Spinal (L) | 196 | 6% |  |  |
|  | Optic Radiations (L) | 109 | 20% |  |  |
| **Component 15** | | **4537** |  | **-36 0 -30** | **9.34** |
| G | Temporal Pole (L) | 1420 | 60% |  |  |
|  | Amygdala (L) | 198 | 56% |  |  |
|  | Hippocampus (L) | 150 | 18% |  |  |
|  | Middle Temporal Gyrus, anterior division (L) | 147 | 33% |  |  |
|  | Pallidum (L) | 137 | 44% |  |  |
|  | Planum Polare (L) | 137 | 38% |  |  |
| W | Inferior Longitudinal Fasciculus (L) | 996 | 58% |  |  |
|  | Uncinate (L) | 372 | 41% |  |  |
|  | Fornix (L) | 189 | 15% |  |  |
|  | Anterior Commissure (L) | 171 | 28% |  |  |
|  | Inferior Occipito Frontal Fasciculus (L) | 156 | 11% |  |  |
|  | Corpus Callosum (L) | 109 | 2% |  |  |
|  | Cortico Spinal (L) | 106 | 3% |  |  |
| **Component 16** | | **5468** |  | **-24 -8 60** | **-6.12** |
| G | Superior Frontal Gyrus (L) | 1678 | 66% |  |  |
|  | Paracingulate Gyrus (L) | 554 | 41% |  |  |
|  | Middle Frontal Gyrus (L) | 274 | 10% |  |  |
|  | Cingulate Gyrus, anterior division (L) | 246 | 25% |  |  |
|  | Juxtapositional Lobule Cortex (Supplementary Motor Cortex) (L) | 198 | 31% |  |  |
|  | Precentral Gyrus (L) | 187 | 4% |  |  |
|  | Cingulate Gyrus, anterior division (R) | 149 | 12% |  |  |
| W | Cingulum (R) | 521 | 12% |  |  |
|  | Corpus Callosum (R) | 497 | 9% |  |  |
|  | Cingulum (L) | 479 | 10% |  |  |
|  | Corpus Callosum (L) | 414 | 7% |  |  |
| **Component 17** | | **2386** |  | **-62 -18 24** | **10.51** |
| G | Postcentral Gyrus (L) | 881 | 24% |  |  |
|  | Precentral Gyrus (L) | 688 | 16% |  |  |
|  | Supramarginal Gyrus, anterior division (L) | 376 | 40% |  |  |
|  | Central Opercular Cortex (L) | 222 | 23% |  |  |
|  | Inferior Frontal Gyrus, pars opercularis (L) | 105 | 14% |  |  |
| **Component 18** | | **1461** |  | **34 -44 22** | **5.89** |
| W | Corpus Callosum (R) | 257 | 5% |  |  |
|  | Arcuate Posterior Segment (R) | 243 | 23% |  |  |
|  | Internal Capsule (R) | 240 | 11% |  |  |
|  | Cortico Spinal (R) | 214 | 8% |  |  |
|  | Inferior Longitudinal Fasciculus (R) | 126 | 7% |  |  |
|  | Cingulum (R) | 122 | 3% |  |  |
|  | Optic Radiations (R) | 120 | 26% |  |  |
| **Component 19 Cluster 1** | | **941** |  | **-22 -74 16** | **5.36** |
| G | Lateral Occipital Cortex, superior division (L) | 144 | 3% |  |  |
| W | Corpus Callosum (L) | 274 | 5% |  |  |
|  | Inferior Longitudinal Fasciculus (L) | 200 | 12% |  |  |
| **Component 19 Cluster 2** | | **890** |  | **-22 -40 26** | **-6.31** |
| W | Cortico Spinal (L) | 255 | 7% |  |  |
|  | Arcuate Posterior Segment (L) | 181 | 24% |  |  |
|  | Corpus Callosum (L) | 116 | 2% |  |  |
| **Component 20 Cluster 1** | | **1076** |  | **-20 -30 44** | **-6.02** |
| W | Cortico Spinal (L) | 272 | 8% |  |  |
|  | Cingulum (L) | 182 | 4% |  |  |
|  | Corpus Callosum (L) | 148 | 3% |  |  |
| **Component 20 Cluster 2** | | **990** |  | **-30 -68 12** | **6.88** |
| W | Inferior Longitudinal Fasciculus (L) | 257 | 15% |  |  |
|  | Corpus Callosum (L) | 211 | 4% |  |  |
|  | Optic Radiations (L) | 128 | 23% |  |  |
| **Component 20 Cluster 3** | | **778** |  | **-28 4 36** | **5.84** |
| W | Corpus Callosum (L) | 182 | 3% |  |  |
|  | Cortico Spinal (L) | 134 | 4% |  |  |

G: grey matter; W: white matter. Coefficents of each component were z-transformed and threshold at |*Z|* > 3.29 (p < 0.001). Clusters larger than 500 voxels were reported. For each cluster, detail regions were reported if it overlapped with more than 100 voxels of a specific region or white matter tract on the templates. We adopted the Harvard-Oxford cortical and subcortical templates, and natbrainlab white matter template. Percentage = significant voxels of this region / total voxels of this region based on the template.
