## Supplementary Table 3 for "Evaluating the granularity and statistical structure of lesions and behaviour in post-stroke aphasia"

**Table 3. Voxel based correlation methodology with behaviours factors region reports**

| Clusters for Behaviour Factors | | Size  (voxels) | Percent | Peak Coordinates  (MINI x, y, z) | Peak  (*r*) |
| --- | --- | --- | --- | --- | --- |
| Phonology | |  |  |  |  |
|  | **Cluster 1** | **5641** |  | **-64 -18 -12** | **0.55** |
| G | Superior Temporal Gyrus, posterior division (L) | 637 | 73% |  |  |
|  | Middle Temporal Gyrus, posterior division (L) | 493 | 39% |  |  |
|  | Supramarginal Gyrus, posterior division (L) | 439 | 42% |  |  |
|  | Planum Temporale (L) | 296 | 56% |  |  |
|  | Planum Polare (L) | 236 | 66% |  |  |
|  | Middle Temporal Gyrus, anterior division (L) | 181 | 41% |  |  |
|  | Heschls Gyrus (includes H1 and H2) (L) | 177 | 55% |  |  |
|  | Insular Cortex (L) | 120 | 10% |  |  |
|  | Central Opercular Cortex (L) | 93 | 9% |  |  |
|  | Middle Temporal Gyrus, temporooccipital part (L) | 86 | 10% |  |  |
|  | Angular Gyrus (L) | 65 | 7% |  |  |
|  | Parietal Operculum Cortex (L) | 46 | 8% |  |  |
|  | Superior Temporal Gyrus, anterior division (L) | 35 | 14% |  |  |
| W | Inferior Longitudinal Fasciculus (L) | 634 | 37% |  |  |
|  | Arcuate Posterior Segment (L) | 547 | 74% |  |  |
|  | Optic Radiations (L) | 198 | 36% |  |  |
|  | Inferior Occipito Frontal Fasciculus (L) | 124 | 9% |  |  |
|  | Cortico Spinal (L) | 114 | 3% |  |  |
|  | Long Segment (L) | 88 | 25% |  |  |
|  | Internal Capsule (L) | 65 | 5% |  |  |
|  | Corpus Callosum (L) | 59 | 1% |  |  |
|  | Arcuate Anterior Segment (L) | 53 | 10% |  |  |
|  | Fornix (L) | 47 | 4% |  |  |
|  | Cortico Ponto Cerebellum (L) | 28 | 3% |  |  |
| Semantics | |  |  |  |  |
|  | **Cluster 1** | **1175** |  | **-36 -12 -18** | **0.54** |
| G | Hippocampus (L) | 133 | 16% |  |  |
|  | Amygdala (L) | 74 | 21% |  |  |
|  | Temporal Pole (L) | 32 | 1% |  |  |
|  | Temporal Fusiform Cortex, posterior division (L) | 30 | 3% |  |  |
|  | Middle Temporal Gyrus, anterior division (L) | 26 | 6% |  |  |
| W | Inferior Longitudinal Fasciculus (L) | 454 | 26% |  |  |
|  | Fornix (L) | 130 | 11% |  |  |
|  | Uncinate (L) | 104 | 11% |  |  |
|  | Inferior Occipito Frontal Fasciculus (L) | 88 | 6% |  |  |
|  | Anterior Commissure (L) | 42 | 7% |  |  |
|  | Optic Radiations (L) | 37 | 7% |  |  |
|  | **Cluster 2** | **509** |  | **-34 -66 18** | **0.46** |
| G | Lateral Occipital Cortex, superior division (L) | 83 | 2% |  |  |
| W | Inferior Longitudinal Fasciculus (L) | 169 | 10% |  |  |
|  | Optic Radiations (L) | 109 | 20% |  |  |
|  | Cortico Spinal (L) | 27 | 1% |  |  |
| Fluency | |  |  |  |  |
|  | **Cluster 1** | **9479** |  | **-60 -6 20** | **0.57** |
| G | Precentral Gyrus (L) | 1537 | 36% |  |  |
|  | Postcentral Gyrus (L) | 786 | 22% |  |  |
|  | Thalamus (L) | 542 | 39% |  |  |
|  | Central Opercular Cortex (L) | 342 | 35% |  |  |
|  | Paracingulate Gyrus (L) | 282 | 21% |  |  |
|  | Caudate (L) | 192 | 34% |  |  |
|  | Brain-Stem (L) | 188 | 9% |  |  |
|  | Cingulate Gyrus, anterior division (L) | 136 | 14% |  |  |
|  | Middle Frontal Gyrus (L) | 111 | 4% |  |  |
|  | Juxtapositional Lobule Cortex (Supplementary Motor Cortex) (L) | 66 | 10% |  |  |
|  | Pallidum (L) | 56 | 18% |  |  |
|  | Hippocampus (L) | 48 | 6% |  |  |
|  | Putamen (L) | 32 | 3% |  |  |
|  | Frontal Pole (L) | 31 | 0% |  |  |
|  | Inferior Frontal Gyrus, pars opercularis (L) | 31 | 4% |  |  |
|  | Cingulate Gyrus, posterior division (L) | 23 | 2% |  |  |
| W | Cortico Spinal (L) | 1444 | 41% |  |  |
|  | Corpus Callosum (L) | 839 | 15% |  |  |
|  | Cingulum (L) | 674 | 14% |  |  |
|  | Internal Capsule (L) | 453 | 37% |  |  |
|  | Arcuate Anterior Segment (L) | 292 | 54% |  |  |
|  | Long Segment (L) | 197 | 56% |  |  |
|  | Fornix (L) | 176 | 14% |  |  |
|  | Superior Cerebelar Pedunculus (L) | 66 | 9% |  |  |
|  | Cortico Ponto Cerebellum (L) | 55 | 7% |  |  |
|  | Anterior Commissure (L) | 52 | 9% |  |  |

G: grey matter; W: white matter. Clusters were significant at Alphasim corrected *p* < 0.01 with voxel *p* < 0.001. For each cluster, detail regions were reported if it overlapped with more than 20 voxels of a specific region or white matter tract on the templates. We adopted the Harvard-Oxford cortical and subcortical templates, and natbrainlab white matter template. Percentage = significant voxels of this region / total voxels of this region based on the template.
