## Supplementary Table 4 for "Evaluating the granularity and statistical structure of lesions and behaviour in post-stroke aphasia"

**Table 4. Regression analysis region reports**

| Clusters for Behaviour Factors | | Size  (voxels) | Percent | Peak Coordinates  (MINI x, y, z) | Peak  (*z*) |
| --- | --- | --- | --- | --- | --- |
| Phonology | |  |  |  |  |
|  | **Cluster 1** | **2546** |  | **-40 -42 2** | **8.12** |
| W | Inferior Longitudinal Fasciculus (L) | 589 | 34% |  |  |
|  | Arcuate Posterior Segment (L) | 387 | 52% |  |  |
|  | Optic Radiations (L) | 300 | 55% |  |  |
|  | Cortico Spinal (L) | 243 | 7% |  |  |
|  | Inferior Occipito Frontal Fasciculus (L) | 143 | 10% |  |  |
|  | Internal Capsule (L) | 117 | 9% |  |  |
|  | Corpus Callosum (L) | 84 | 1% |  |  |
|  | Arcuate Anterior Segment (L) | 77 | 14% |  |  |
|  | Long Segment (L) | 76 | 22% |  |  |
|  | Fornix (L) | 68 | 6% |  |  |
|  | Cortico Ponto Cerebellum (L) | 43 | 5% |  |  |
|  | **Cluster 2** | **1393** |  | **-40 32 4** | **6.29** |
| G | Inferior Frontal Gyrus, pars opercularis (L) | 239 | 33% |  |  |
|  | Frontal Operculum Cortex (L) | 156 | 43% |  |  |
|  | Central Opercular Cortex (L) | 131 | 13% |  |  |
|  | Inferior Frontal Gyrus, pars triangularis (L) | 100 | 16% |  |  |
|  | Precentral Gyrus (L) | 90 | 2% |  |  |
|  | Insular Cortex (L) | 22 | 2% |  |  |
| W | Inferior Occipito Frontal Fasciculus (L) | 131 | 10% |  |  |
|  | Uncinate (L) | 59 | 6% |  |  |
| Semantics | |  |  |  |  |
|  | **Cluster 1** | **2577** |  | **-32 -2 -30** | **5.73** |
| G | Temporal Pole (L) | 464 | 20% |  |  |
|  | Amygdala (L) | 156 | 44% |  |  |
|  | Putamen (L) | 132 | 14% |  |  |
|  | Hippocampus (L) | 118 | 14% |  |  |
|  | Pallidum (L) | 102 | 33% |  |  |
|  | Thalamus (L) | 54 | 4% |  |  |
|  | Temporal Fusiform Cortex, posterior division (L) | 42 | 5% |  |  |
|  | Middle Temporal Gyrus, anterior division (L) | 27 | 6% |  |  |
|  | Inferior Temporal Gyrus, anterior division (L) | 23 | 7% |  |  |
| W | Inferior Longitudinal Fasciculus (L) | 651 | 38% |  |  |
|  | Uncinate (L) | 223 | 24% |  |  |
|  | Fornix (L) | 204 | 17% |  |  |
|  | Anterior Commissure (L) | 162 | 27% |  |  |
|  | Inferior Occipito Frontal Fasciculus (L) | 124 | 9% |  |  |
|  | Optic Radiations (L) | 99 | 18% |  |  |
|  | Cortico Spinal (L) | 97 | 3% |  |  |
|  | Internal Capsule (L) | 64 | 5% |  |  |
|  | Superior Cerebelar Pedunculus (L) | 24 | 3% |  |  |
|  | **Cluster 2** | **1117** |  | **-58 -38 40** | **7.18** |
| G | Supramarginal Gyrus, anterior division (L) | 353 | 37% |  |  |
|  | Supramarginal Gyrus, posterior division (L) | 332 | 32% |  |  |
|  | Parietal Operculum Cortex (L) | 210 | 36% |  |  |
|  | Angular Gyrus (L) | 176 | 19% |  |  |
| W | Arcuate Posterior Segment (L) | 35 | 5% |  |  |
|  | **Cluster 3** | **396** |  | **-32 -62 32** | **5.92** |
| W | Optic Radiations (L) | 63 | 11% |  |  |
|  | Corpus Callosum (L) | 53 | 1% |  |  |
|  | Inferior Longitudinal Fasciculus (L) | 41 | 2% |  |  |
| Executive | |  |  |  |  |
|  | **Cluster 1** | **2434** |  | **-44 -78 20** | **5.66** |
| G | Lateral Occipital Cortex, superior division (L) | 636 | 13% |  |  |
|  | Lateral Occipital Cortex, inferior division (L) | 481 | 24% |  |  |
|  | Middle Temporal Gyrus, posterior division (L) | 369 | 29% |  |  |
|  | Inferior Temporal Gyrus, posterior division (L) | 199 | 20% |  |  |
|  | Inferior Temporal Gyrus, anterior division (L) | 167 | 51% |  |  |
|  | Middle Temporal Gyrus, temporooccipital part (L) | 126 | 15% |  |  |
|  | Temporal Pole (L) | 90 | 4% |  |  |
|  | Middle Temporal Gyrus, anterior division (L) | 89 | 20% |  |  |
|  | Inferior Temporal Gyrus, temporooccipital part (L) | 60 | 9% |  |  |
| W | Inferior Longitudinal Fasciculus (L) | 50 | 3% |  |  |
|  | Optic Radiations (L) | 34 | 6% |  |  |
|  | Cortico Spinal (L) | 22 | 1% |  |  |
|  | **Cluster 2** | **600** |  | **-44 34 24** | **6.80** |
| G | Middle Frontal Gyrus (L) | 390 | 14% |  |  |
|  | Frontal Pole (L) | 126 | 2% |  |  |
|  | Inferior Frontal Gyrus, pars triangularis (L) | 75 | 12% |  |  |
| Fluency | |  |  |  |  |
|  | **Cluster 1** | **3306** |  | **-52 -2 40** | **6.87** |
| G | Middle Frontal Gyrus (L) | 1059 | 38% |  |  |
|  | Precentral Gyrus (L) | 1001 | 23% |  |  |
|  | Inferior Frontal Gyrus, pars opercularis (L) | 82 | 11% |  |  |
|  | Postcentral Gyrus (L) | 40 | 1% |  |  |
| W | Corpus Callosum (L) | 184 | 3% |  |  |
|  | Cortico Spinal (L) | 116 | 3% |  |  |
|  | Cingulum (L) | 56 | 1% |  |  |
|  | **Cluster 2** | **2547** |  | **-22 -6 12** | **7.95** |
| G | Putamen (L) | 425 | 46% |  |  |
|  | Thalamus (L) | 336 | 24% |  |  |
|  | Caudate (L) | 175 | 31% |  |  |
|  | Pallidum (L) | 136 | 44% |  |  |
| W | Cortico Spinal (L) | 851 | 24% |  |  |
|  | Internal Capsule (L) | 244 | 20% |  |  |
|  | Inferior Occipito Frontal Fasciculus (L) | 155 | 11% |  |  |
|  | Fornix (L) | 139 | 11% |  |  |
|  | Corpus Callosum (L) | 47 | 1% |  |  |

G: grey matter; W: white matter. Behavioural voxel coefficients = stepwise regression model coefficients × principal component analysis coefficients. Then coefficients were z-transformed and thresholded at |*Z|* > 3.29 (*p* < 0.001). Clusters larger than 200 voxels were reported. For each cluster, detail regions were reported if it overlapped with more than 20 voxels of a specific region or white matter tract on the templates. We adopted the Harvard-Oxford cortical and subcortical templates, and natbrainlab white matter template. Percentage = significant voxels of this region / total voxels of this region based on the template.
